# miR-425-5p Regulates Cellular Senescence Through Modulation of Retinoblastoma Protein Phosphorylation

**DOI:** 10.64898/2026.06.15.732173

**Authors:** Latika Matai, Sarah Haggenmueller, Jonathan D. Lee, Frank J Slack

**Affiliations:** Department of Pathology, Beth Israel Deaconess Medical Center, Harvard Medical School, 330 Brookline Ave, Boston, MA, 02215, United States; Harvard Medical School Initiative for RNA Medicine, Harvard Medical School, Boston, MA, United States; School of Natural Sciences, Department of Bioscience, Technical University Munich, 85748 Garching, Germany

**Author notes:** Co-corresponding Authors Corresponding authors: Latika Matai and Frank J. Slack.

## Abstract

MicroRNAs (miRNAs) are small non-coding RNAs that play critical roles in regulating cellular senescence and aging. Our recent studies identified a conserved *C. elegans* miRNA cluster (miR-229/64/65/66) that is required for normal adult lifespan, with overexpression significantly extending longevity. Notably, cel-miR-229 is evolutionarily conserved in humans, with hsa-miR-425 sharing an identical seed sequence.

Here, we investigated the role of miR-425 in mammalian cellular senescence. We found that miR-425 expression is markedly reduced in pharmacologically induced senescence in human lung cancer cells. Restoration of miR-425 expression attenuates senescence and suppresses the expression of senescence-associated secretory phenotype (SASP) cytokines following senescence induction. We further observed that miR-425 levels decline during replicative senescence, whereas stable overexpression in WI-38 fibroblasts delays senescence accumulation and preserves proliferative capacity.

Mechanistically, miR-425 suppresses TGF-β signaling, leading to reduced expression of the cyclin-dependent kinase inhibitor p21/*CDKN1A* and increased phosphorylation of the retinoblastoma (RB) protein, thereby promoting cell-cycle progression. We further identify PPP2CB, the catalytic subunit of protein phosphatase 2A (PP2A), as a direct target of miR-425. *PPP2CB* expression is downregulated in miR-425-5p overexpressing cells, even under senescence induction. Knockdown of *PPP2CB* using siRNA phenocopies the effects of miR-425 overexpression, reducing senescence, enhancing proliferative potential, and increasing RB phosphorylation.

Collectively, our findings identify miR-425 as a conserved regulator of cellular senescence that acts through upregulation of RB phosphorylation. These results establish a novel miR-425-*PPP2CB*-RB regulatory axis controlling proliferation and senescence and suggest miR-425 as a potential therapeutic target for mitigating senescence to promote extended health span.

## Introduction

Cellular senescence is a fundamental hallmark of aging characterized by a stable cell-cycle arrest, profound changes in gene expression and metabolism, and the acquisition of a pro-inflammatory senescence-associated secretory phenotype (SASP)^1–4^. While transient senescence can play beneficial roles in tissue repair and tumor suppression, the progressive accumulation of senescent cells with age contributes to chronic inflammation, impaired tissue regeneration, functional decline, and the development of numerous age-related diseases, including neurodegenerative, cardiovascular, and metabolic disorders^5–11^. Importantly, genetic, and pharmacological interventions that reduce senescent cell burden or suppress senescence pathways have been shown to improve tissue function, restore tissue homeostasis, extend health-span, and delay age-related pathologies in animal models^12^. For example, clearance of p16INK4a-positive senescent cells improves physical function and lifespan in mice, while caloric restriction delays cellular senescence and preserves tissue function^13–15^. Consequently, there is growing interest in identifying molecular regulators of senescence and the mechanisms through which they influence cellular aging^16,17^. Elucidating these regulatory mechanisms may uncover novel therapeutic targets to attenuate senescence burden, restore tissue homeostasis, mitigate age-associated functional decline, and ultimately extend health-span.

Cellular senescence can be broadly categorized into replicative senescence and stress-induced senescence. Replicative senescence arises as a consequence of repeated cell division and progressive telomere shortening and is considered a physiologically relevant model of organismal aging and age associated tissue decline^18^. In contrast, therapy- or drug-induced senescence is triggered by cellular stressors such as chemotherapy, radiation, or oxidative damage and is characterized by the rapid activation of senescence pathways^19^. Drug induced senescence is increasingly recognized as a contributor to chronic inflammation, tissue dysfunction, and accelerated aging in cancer survivors^20^. Therefore, studying both forms of senescence is important, as it enables the identification of conserved regulators of senescence while also revealing pathways that may be specifically relevant to aging or therapeutic interventions. At the molecular level, senescence is primarily enforced through activation of the cyclin-dependent kinase (CDK) inhibitors p21 and p16^INK4a^, which inhibits CDKs and prevent the phosphorylation of the retinoblastoma protein (RB)^21^. Hypo-phosphorylated RB remains active and represses E2F-dependent transcription, thereby blocking cell cycle progression and establishing the senescent growth arrest^22^. RB thus serves as a key central effector of senescence and an important determinant of whether cells continue to proliferate or enter a stable senescent state^23^.

MicroRNAs (miRNAs) are a class of small (∼22 nucleotide) non-coding RNAs that regulate gene expression post-transcriptionally by binding to complementary sequences within target mRNAs, leading to mRNA degradation or translational repression^24^. Increasing evidence has established miRNAs play an intertwined role with senescence, driving age-related complications^25–29^. MiRNAs are differentially expressed during senescence, sometimes regulated by SASP factors^30–32^. Conversely, several miRNAs, including miR-34a, miR-217, and miR-146a, have been shown to promote senescence by regulating p53, p21, inflammatory signaling, and DNA damage response pathways, whereas others, such as members of the *let-7* family and miR-17-92 cluster, have been implicated in maintaining cellular homeostasis and delaying age-associated decline^32–37^. Due to ease of synthesis and manipulation, miRNAs represent attractive therapeutic candidates for modulating senescence and promoting healthy aging^38,39^. Given the central role of miRNAs in aging and senescence, identifying evolutionarily conserved miRNA regulators may provide insight into fundamental mechanisms governing cellular aging across species.

*C. elegans* has been instrumental in identifying conserved regulators of aging, including longevity-associated miRNAs. Many aging-related miRNAs first identified in *C. elegans*, have proven to be remarkably conserved across evolution^40,41^. In our previous work, we demonstrated that the *C. elegans* miR-229/64/65/66 cluster promotes longevity^42^. Notably, hsa-miR-425 shares an identical seed sequence with cel-miR-229, suggesting conservation of target recognition and biological function across species. Given this evolutionary relationship, we have now examined the role of miR-425 in mammalian systems.

Previous studies have implicated miR-425 in the regulation of cellular proliferation, apoptosis, and stress responses across diverse biological contexts. Depending on the cellular environment, miR-425 has been reported to exert both pro-survival and growth-suppressive effects, reflecting the diverse signaling pathways under its control. In several cancer models, miR-425 promotes proliferation, migration, and survival through repression of *PTEN* and activation of the PI3K/AKT pathway^43^. Conversely, other studies have demonstrated that miR-425 suppresses proliferation and induces apoptosis through regulation of targets such as *E2F6* and *PAK4*^44,45^. These observations suggest that miR-425 functions as a concentration and context-dependent regulator of cellular homeostasis and cell-fate decisions.

Despite growing evidence linking miR-425 to the regulation of proliferation, apoptosis, and stress-response pathways, its role in cellular senescence remains unknown. Here, we investigated whether miR-425-5p regulates both therapy-induced and replicative senescence and explored the underlying molecular mechanism. We show that miR-425-5p is consistently downregulated during senescence and that its restoration suppresses senescence-associated phenotypes, including SASP induction. Furthermore, our findings suggest that regulation of RB phosphorylation is an important component of a miR-425-5p-regulated pathway controlling cellular senescence.

## Results

### miR-425-5p overexpression (OE) mitigates drug induced senescence

Our previous work identified cel-miR-229/64/65/66 cluster as a regulator of longevity in *C. elegans,* where its overexpression extends lifespan and deletion of the miRNA cluster shortens lifespan^42^. cel-miR-229 is highly conserved, with several human homologs, including hsa-miR-425, which shares an identical seed sequence (Fig. S1a). We therefore sought to investigate the role of hsa-miR-425-5p (miR-425-5p hereafter) in regulating cellular senescence. To evaluate its function across distinct senescence contexts, we utilized both drug induced and replicative senescence models. For drug induced senescence, we employed methotrexate (MTX), a widely used chemotherapeutic and immunomodulatory agent that induces senescence through nucleotide depletion, causing DNA damage and oxidative stress leading to activation of cell-cycle arrest pathways. MTX-induced senescence is a clinically relevant model because therapy-induced senescent cells can persist in tissues, promote chronic inflammation through SASP factors, and contribute to long-term tissue dysfunction and accelerated aging^46^. Consequently, MTX-induced senescence provides a clinically relevant and experimentally tractable model to investigate the molecular mechanisms underlying senescence and to identify interventions that mitigate senescent cell accumulation while preserving therapeutic efficacy. Additionally, drug induced senescence shares many molecular and phenotypic features with naturally occurring age-associated senescence, including activation of p53/p21 signaling, persistent DNA damage responses, SASP production, and loss of viability^47^.

To investigate whether miR-425-5p is regulated during drug induced senescence, A549 cells were treated with methotrexate (MTX), and mature miR-425-5p levels were quantified. miR-425-5p expression was significantly reduced following MTX treatment compared with untreated controls (Fig. 1a), suggesting a potential role in senescence regulation. To determine whether restoration of miR-425-5p could attenuate senescence, A549 cells were transfected with a moderate amount of miR-425-5p or negative control mimic for 36 h, followed by treatment with methotrexate (MTX) to induce senescence (Fig. 1b). Transfection with mimic leads to successful sustained upregulation of mature miR-425-5p levels in cells, to the timepoint when all the readouts were performed (Fig S1b). Senescence was assessed by lysosomal β-galactosidase activity using a blue/white cell imaging assay. Importantly, restoration of miR-425-5p expression prior to MTX exposure significantly decreased the proportion of senescent cells relative to negative control-transfected cells (Fig. 1c,d), suggesting a protective role for miR-425-5p against drug induced cellular senescence. Consistent with this previous literature, miR-425-5p overexpression also enhanced the proliferative capacity of A549 cells (Fig. S1c).

**Figure 1:**
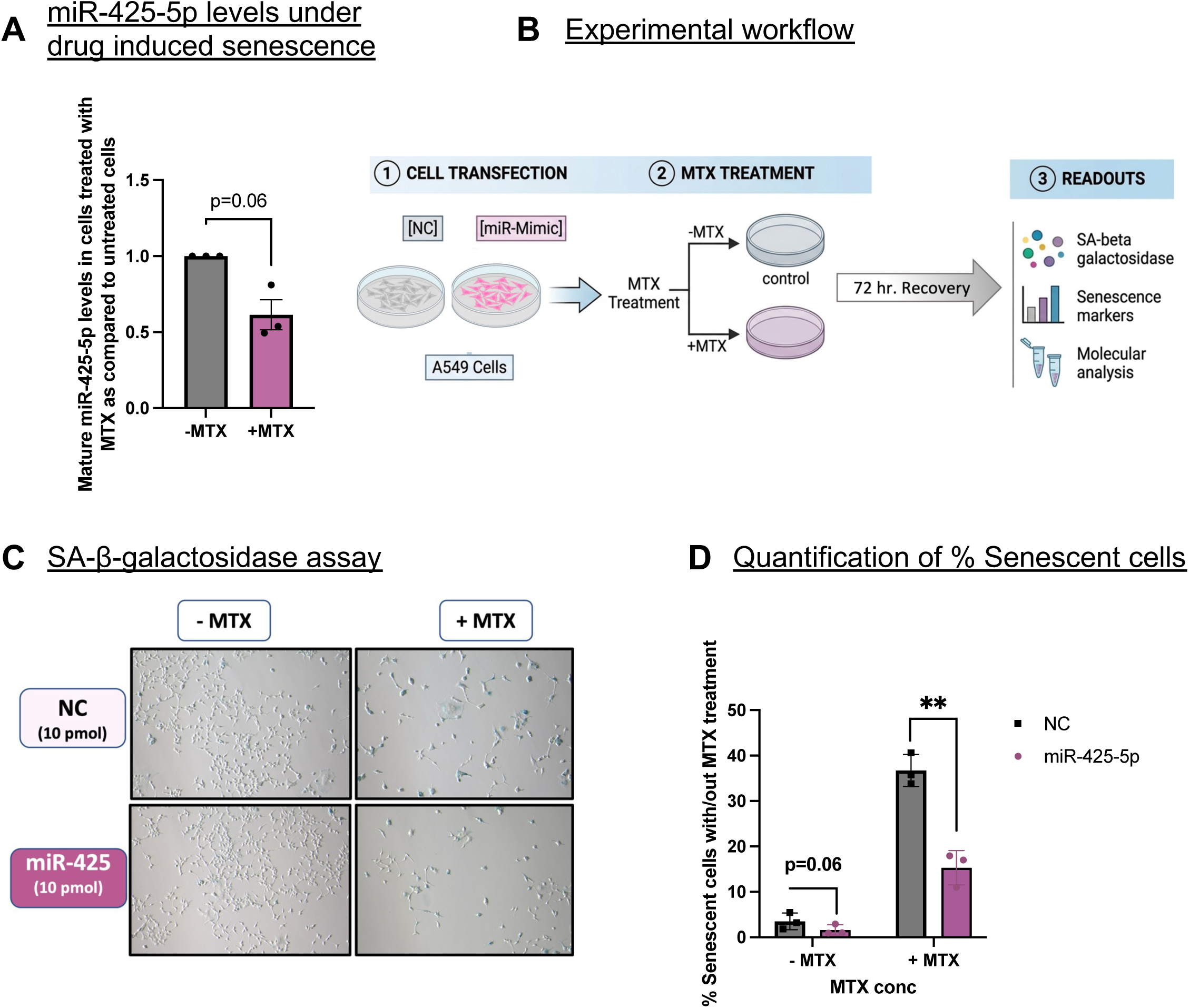
miR-425-5p overexpression reduces drug induced senescence. A) The graph represents relative expression of mature miR-425-5p in A549 cells treated with 8 μM methotrexate (MTX) for 24 hours, followed by a 72-hour recovery period, compared with untreated cells collected at the same time point, indicating that miR-425-5p is downregulated during MTX-induced cellular senescence. * indicates p<0.05, as. determined by a student’s t-test. B) Experimental workflow for assessing the effect of miR-425-5p on methotrexate (MTX)-induced cellular senescence in A549 cells. A549 cells were transfected with either a negative control (NC) mimic or miR-425-5p mimic. Thirty-six hours post-transfection, transfected cells were split and either left untreated or exposed to methotrexate (MTX 8µM) to induce senescence. Following a 24-hour MTX treatment, the drug-containing medium was removed, and cells were washed with PBS and allowed to recover in fresh growth medium for 72 hours. Senescence-associated phenotypes were subsequently evaluated by SA-β-galactosidase staining, analysis of senescence markers, and molecular characterization by miRNA, RNA and protein-based assays. C) Representative images of cells stained for senescence-associated β-galactosidase (SA-β-gal) activity following a 72-hour recovery period after MTX treatment/untreated in cells previously transfected with either a negative control mimic or miR-425-5p mimic. Representative images demonstrate a marked reduction in the percentage of SA-β-gal-positive blue cells in miR-425-5p-overexpressing cells following MTX treatment, indicating that miR-425-5p attenuates MTX-induced cellular senescence. D) Quantification of percent senescent (blue) cells from three independent biological replicates. miR-425-5p overexpression reduced basal senescence and significantly attenuated MTX-induced senescence, as indicated by a lower percentage of SA-β-gal-positive blue cells compared with negative control-transfected cells.

In contrast, inhibition of miR-425-5p using a specific antagomir resulted in efficient knockdown of mature miR-425-5p levels (Fig. S1e) and slightly increases the proportion of senescent cells following MTX treatment (Fig. S1d). Interestingly, the protective effects of miR-425-5p (mimic) were observed only at moderate levels of overexpression. Higher miR-425-5p mimic concentrations increased apoptosis, as measured by Annexin V staining (Fig. S1f) and failed to confer protection against senescence (Fig. S1g), suggesting that miR-425-5p exerts a dose-dependent, biphasic effect on cellular senescence where lower concentration of the miRNA is protective towards senescence, unlike the higher, detrimental concentration.

### MiR-425-5p overexpression delays replicative senescence

We also sought to test the role of miR-425-5p in replicative/proliferative senescence. For this we choose to work on WI-38 human diploid lung fibroblasts, which are one of the most extensively characterized and widely used models of replicative senescence. Unlike transformed or immortalized cell lines, WI-38 cells, during serial passaging, ultimately enter a stable senescent state that closely recapitulates physiological aging. WI-38 cells exhibit hallmark features of cellular senescence at higher population doubling levels (PDL 58-62), including irreversible cell-cycle arrest and increased SA-β-gal activity. Consistent with our findings in drug induced senescence, miR-425-5p expression appears to progressively decline with the onset of replicative senescence in WI-38 cells (Fig. S2a). To investigate its functional role, we generated stable miR-425-5p-overexpressing WI-38 cells using a lentiviral vector (AbmGood) alongside a scrambled control (Fig. S2b). Notably, miR-425-5p overexpression delayed the accumulation of senescent cells compared with control population (Fig. 2a,b). In addition, miR-425-5p-overexpressing cells exhibited enhanced proliferative capacity, as demonstrated by increased MTT activity at higher population doubling levels (PDLs), when control cells showed a marked decline in proliferative potential (Fig. 2c). Together, these findings indicate that miR-425-5p delays replicative senescence and promotes proliferative potential in WI-38 fibroblasts.

**Figure 2:**
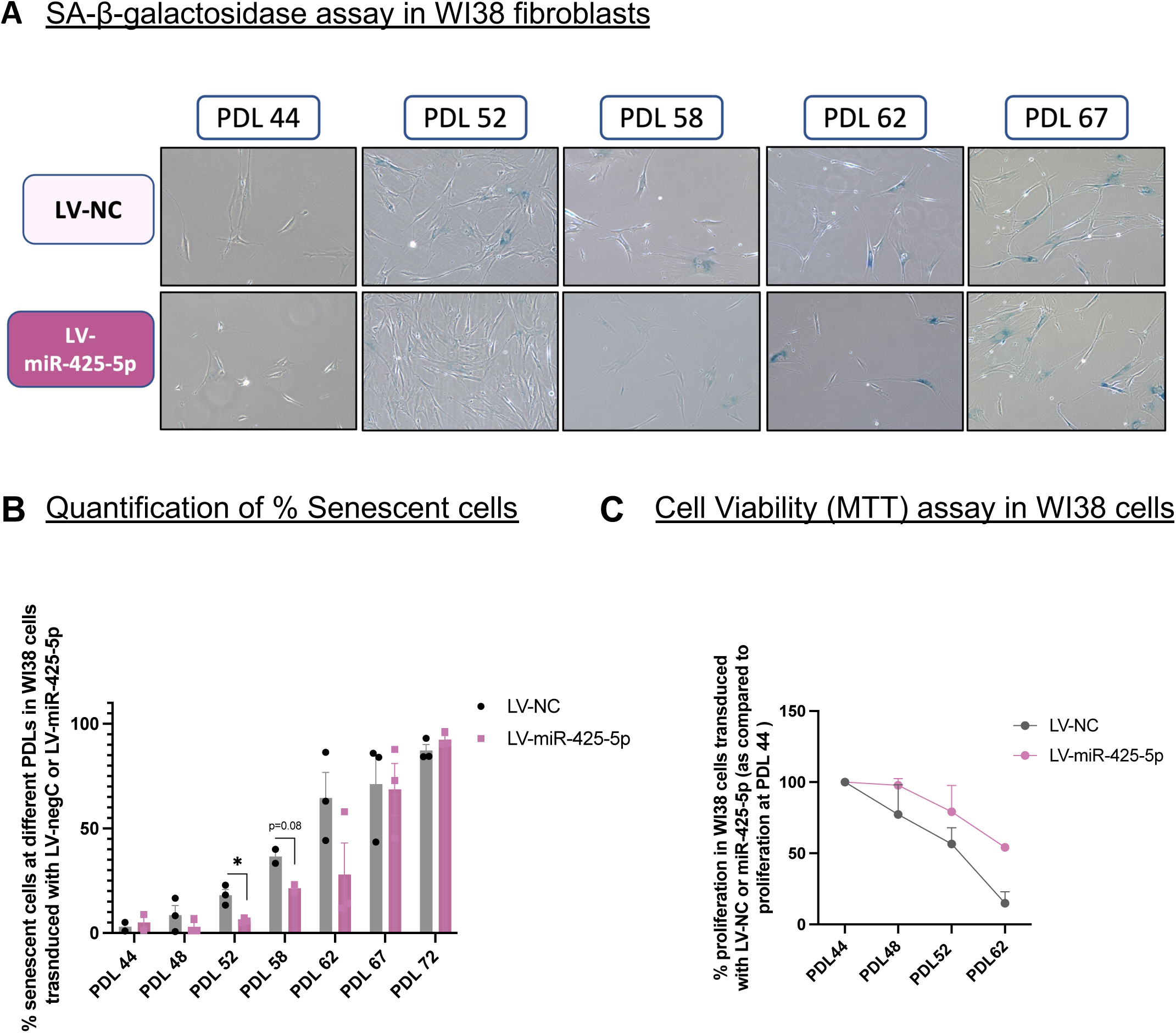
miR-425-5p overexpression delays replicative senescence. A) Representative images of stable miR-425-5p-overexpressing and negative control WI-38 fibroblasts stained for senescence-associated β-galactosidase (SA-β-gal) activity at the indicated population doubling levels (PDLs). Representative images show a marked reduction in blue cells in miR-425-5p-overexpressing cultures at PDL 52 and PDL 58, indicating that miR-425-5p delays the accumulation of replicative senescence. B) Quantification of the percentage of senescent (blue) WI-38 cells from three independent biological replicates at the indicated population doubling levels (PDLs), demonstrating delayed accumulation of replicative senescence upon miR-425-5p overexpression. C) Percentage cell proliferation in stable miR-425-5p-overexpressing and negative control WI-38 fibroblasts at the indicated population doubling levels (PDLs), compared to PDL 44 as measured by the CyQUANT MTT assay. miR-425-5p-overexpressing cells exhibited increased proliferative capacity compared with control cells, particularly at later PDLs.

### miR-425-5p OE decreases Senescence Associated Secretory Phenotype (SASP) induction

A key feature of senescent cells is the senescence-associated secretory phenotype (SASP), which consists of inflammatory cytokines, chemokines, growth factors, and other molecules secreted by senescent cells. These factors can promote chronic inflammation, impair tissue function, and induce senescence in neighboring cells. Therefore, measuring SASP factors is commonly used to assess the extent of senescence and the effectiveness of interventions that suppress senescence. Common SASP markers include the inflammatory cytokines IL-6, IL-8, as well as chemokines such as CCL2 (MCP-1) and CXCL1. To determine whether miR-425-5p regulates the senescence-associated secretory phenotype (SASP), we measured the expression of selected cytokines and chemokines following senescence induction. miR-425-5p overexpression significantly reduced the expression of most SASP-associated factors in A549 cells, both under basal conditions and following senescence induction (Fig. 3). In contrast, no significant differences were observed in DNA damage signaling between control and miR-425-5p-overexpressing cells, as assessed by γ-H2AX levels normalized to total H2AX following MTX treatment (Fig. S3a). We also observed a modest reduction in phosphorylated ATM levels in miR-425-5p-overexpressing cells under basal conditions; however, this effect was not maintained following senescence induction (Fig. S3b).

**Figure 3:**
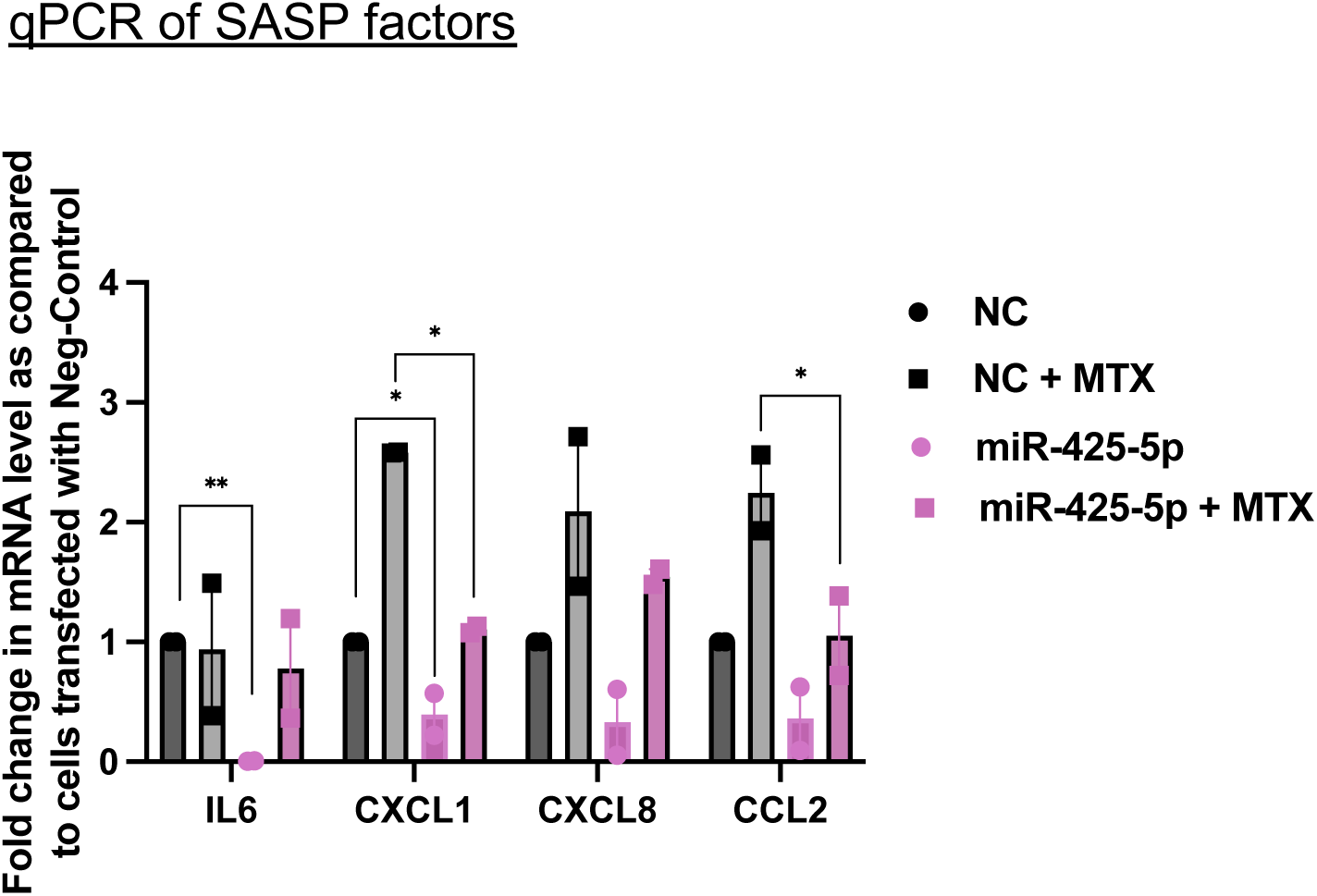
miR-425-5p overexpression attenuates Senescence-Associated Secretory Phenotype (SASP) induction. Graph showing fold changes in mRNA expression of SASP-associated cytokines in neg-control A549 cells treated with MTX, miR-425-5p OE cells, untreated and treated with MTX, as compared to untreated negative control cells (NC). Error bars show SEM, *indicates p<0.05, as. determined by a student’s t-test. miR-425-5p overexpression reduced the induction of multiple SASP factors following senescence induction.

### miR-425 overexpression downregulates the TGF-β/*CDKN1A* axis and enhance RB phosphorylation

In order to investigate the target genes and pathways regulated by miR-425-5p, we analyzed miR-425-5p predicted targets via TargetScan using DAVID gene ontology analysis (https://davidbioinformatics.nih.gov/) (Fig 4a). “Cellular senescence” pathway was found enriched in miR-425 predicted targets (Fig 4a). Most of these targets were found downregulated in miR-425-5p OE cells, suggesting potential interaction and regulation by the miRNA (Fig. S4a). Noteworthy, miR-425-5p has potential targets in TGF-Beta pathways including *TGFB1* (ligand), *TGFBR2* (the receptor) and *SMAD2* (the effector SMAD) (Fig 4a). We found that these levels of these transcripts are significantly decreased in miR-425 OE cells (Fig 4b).

**Figure 4:**
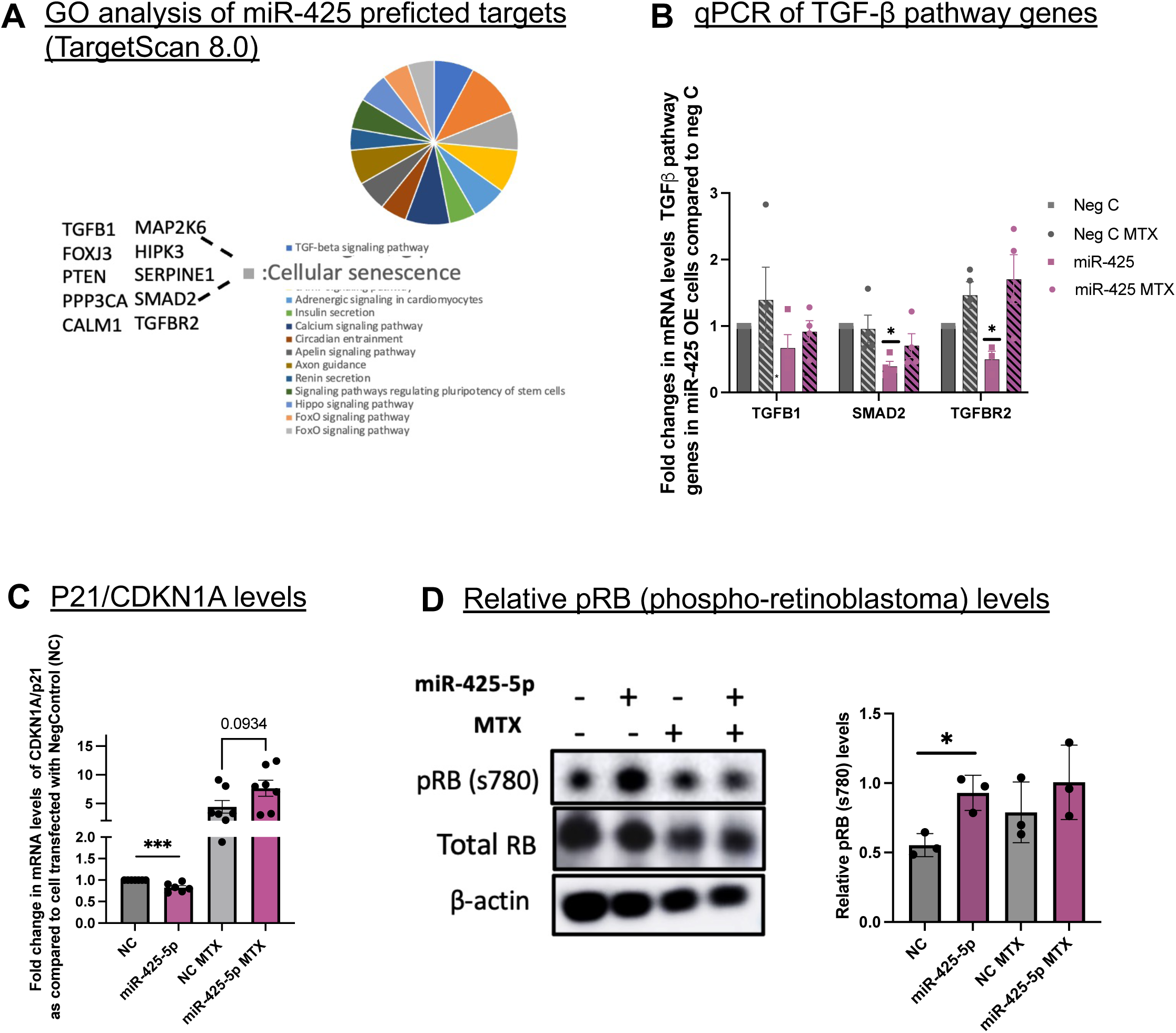
miR-425 overexpression downregulates the TGF-β/CDKN1A axis and enhance RB Phosphorylation. A) Chart showing GO/KEGG pathway enrichment analysis of miR-425-5p predicted targets using TargetScan database, indicating that cellular senescence pathway is enriched in miR-425-5p regulated transcripts. B) Graph represents fold changes in mRNA levels of *TGFB1, SMAD2* and *TGFBR2* in neg-control A549 cells treated with MTX, miR-425-5p OE cells, untreated and treated with MTX, as compared to untreated negative control cells (NC). Error bars show SEM, *indicates p<0.05, as. determined by a student’s t-test. This indicates downregulation of TGF-beta pathways transcripts in miR-425-5p OE cells in untreated and senescence inducing conditions. C) Graph represents fold changes in mRNA levels of *CDKN1A/p21* cell cycle arrest gene in neg-control A549 cells treated with MTX, miR-425-5p OE cells, untreated and treated with MTX, as compared to untreated negative control cells (NC). Error bars show SEM, *indicates p<0.05, as. determined by a student’s t-test. This indicates downregulation of *CDKN1A* transcript in miR-425-5p OE cells in untreated condition. D) Representative western blots and corresponding quantification of phosphorylated RB (Ser780), total RB, and β-actin levels in negative control and miR-425-5p-overexpressing A549 cells under untreated and MTX-induced senescence conditions. The graph on the right shows quantification of pRB (Ser780) levels normalized to total RB, with total RB levels normalized to the loading control β-actin. Data represent three independent biological replicates. *indicates p<0.05, as. determined by a student’s t-test.

TGF-β is a well-established inducer of cell-cycle arrest and senescence through activation of canonical SMAD signaling^48^. Upon TGF-β receptor activation, SMAD2/3 form complexes with SMAD4 and translocate to the nucleus, where they transcriptionally induce cyclin-dependent kinase inhibitors (CDKIs), most notably p15^INK4B^ (*CDKN2B*) and p21^CIP1^ (*CDKN1A*)^49–51^. These CDKIs inhibit Cyclin D–CDK4/6 and Cyclin E–CDK2 complexes, thereby preventing phosphorylation of the retinoblastoma protein (RB)^51–53^. Hypophosphorylated RB binds and represses E2F transcription factors, resulting in G1 arrest and senescence-associated growth inhibition^54–56^. In A549 cells, however, *CDKN2B* and *CDKN2A* (p15^INK4B^ and p16^INK4A^) are deleted due to loss of the 9p21 locus, making p21^CIP1^ the predominant mediator of TGF-β-induced cell-cycle arrest. Consequently, likely due to the suppression of TGF-β/SMAD signaling, we observed reduced p21 expression in miR-425-5p OE cells (Fig. 4c), whereas this reduction was not observed in MTX-treated cells, probably due to strong effects of MTX on senescence shadowing the protective effects of miRNA (Fig. 4c). We also observed increased cyclin D1 expression in miR-425-5p OE cells (Fig. S4c), consistent with previous findings that miR-425 stimulates the PI3K/AKT pathway and thereby promotes cyclin D1 expression^43^. We hypothesize that this increase in cyclin D1 levels could also result from the regulation of TGF-β signaling and inhibition of CDK inhibitor p21^57^. Since TGF-β regulation of p21 is a p53-independent mechanism, we observed no reduction, and in fact an increase, in p53 levels in miR-425-5p OE cells despite the suppression of senescence^51^ (Fig. S4b).

Most importantly, we observed increased phosphorylation of retinoblastoma protein, pRB (Ser780), in miR-425-5p OE cells (Fig. 4d). Increased phosphorylation of RB at Ser780 is consistent with reduced RB-mediated cell-cycle repression and may contribute to the attenuation of senescence observed in miR-425-overexpressing cells (Figure 1c,d). Since maintenance of RB in a hypophosphorylated state is critical for establishment of senescence-associated growth arrest, increased pRB levels suggest enhanced proliferative potential and diminished senescence signaling^58^. The RB phosphorylation appears to be upregulated in miR-425-5p OE cells with MTX treatment (Fig 4d) suggesting additional factors, other than p21 inhibition playing a role in this sustained upregulation.

### miR-425-5p targets *PPP2CB*, the catalytic subunit of PP2A, to regulate RB phosphorylation and cellular senescence

To identify potential miR-425-5p targets involved in the regulation of cellular senescence, we performed RNA-sequencing analysis in A549 cells transfected with either a negative control mimic or miR-425-5p mimic under both untreated and senescence-inducing conditions (Fig 1b). A total of 151 transcripts were significantly downregulated in miR-425-5p-overexpressing cells compared with control cells under basal conditions (Fig. S5a). GO and KEGG pathway enrichment analyses revealed significant enrichment of pathways associated with longevity regulation, complement activation, MAPK signaling, autophagy, and NOD-like receptor signaling (Fig. S5a). Following senescence induction, the transcriptional differences between miR-425-5p-overexpressing and control cells were less pronounced, likely reflecting the dominant effects of MTX-induced DNA damage on global gene expression. Nevertheless, pathways related to complement activation, autophagy, and cellular metabolism remained significantly enriched among differentially expressed genes (Fig. S5b).

To identify potential direct targets of miR-425-5p, we compared transcripts that were consistently downregulated in miR-425-5p OE cells, both in the presence and absence of senescence induction, with predicted miR-425 targets identified through TargetScan (Fig. 5a). This analysis identified two candidate targets, *GLIS3* and *PPP2CB*, the latter encoding the catalytic subunit of PP2A phosphatase (Fig. 5a)^59,60^. PP2A is known to dephosphorylate several proteins, including RB, and other pocket proteins p107 and p130^61–64^. Although the other pocket proteins are considered major PP2A substrates, PP2A has also been reported to dephosphorylate RB under conditions of oxidative stress, such as those induced by MTX treatment^65^.

**Figure 5:**
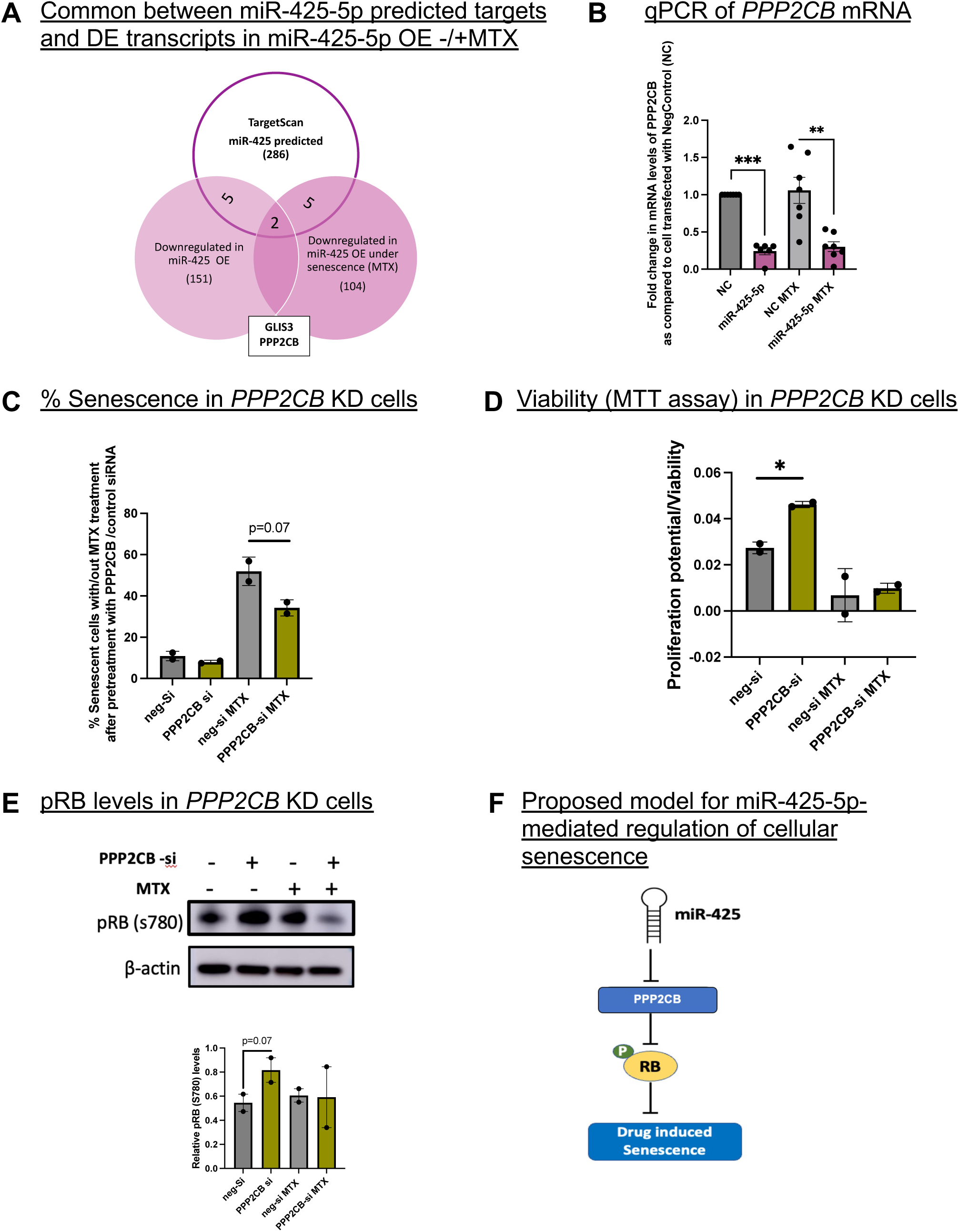
miR-425-5p Targets *PPP2CB* (encoding the Catalytic Subunit of PP2A), to Regulate RB Phosphorylation and Cellular Senescence. A) Pie chart representing transcripts that are common between miR-425-5p predicted targets identified using TargetScan database and transcripts that were downregulated in miR-425-5p OE A549 cells in basal and senescence inducing conditions, identifying *GLIS3* and *PPP2CB*. B) Graph represents fold changes in mRNA levels of *PPP2CB* gene in Neg-control A549 cells treated with MTX, miR-425-5p OE cells, untreated and treated with MTX, as compared to untreated negative control cells (NC). Error bars show SEM, *indicates p<0.05, as. determined by a student’s t-test. This indicates significant downregulation of *PPP2CB* transcript in miR-425-5p OE cells in basal condition and with MTX treatment. C) Graph represents quantification of percent senescent (blue) cells after MTX treatment/untreated in cells transfected with either a negative control siRNA or *PPP2CB* siRNA. This indicates marked reduction in the percentage of senescent cells under *PPP2CB* knockdown. D) Quantification of cell proliferation in *PPP2CB* siRNA and negative control siRNA treated A549 cells, with and without MTX induction, as measured by the CyQUANT MTT assay. *PPP2CB* KD cells exhibited increased proliferative capacity compared with control cells. E) Representative western blots and corresponding quantification of phosphorylated RB (Ser780) and β-actin levels in negative control and *PPP2CB* siRNA treated A549 cells under untreated and MTX-induced senescence. The graph on the right shows quantification of pRB (Ser780) levels normalized to the loading control β-actin. F) Model representing miR-425-5p regulation of senescence through modulation of RB phosphorylation by targeting *PPP2CB* mRNA.

miR-425-5p is predicted to bind the *PPP2CB* 3′UTR at two sites, one of which displays features of a canonical 7mer-A1 binding site (Fig. S5c). We observed significant downregulation of *PPP2CB* transcripts in miR-425-5p OE cells under both basal and senescence-inducing conditions, as identified by qPCR, thereby validating our RNA-seq findings (Fig. 5b). To test whether *PPP2CB* is a key mediator of miR-425-5p-regulated senescence, we performed siRNA-mediated knockdown of *PPP2CB* in A549 cells. Knockdown of *PPP2CB* reproduced the biological effects observed with miR-425-5p overexpression, including suppression of senescence, and enhanced cellular proliferation in A549 cells (Fig. 5c,d). Consistent with these findings, *PPP2CB* transcript levels robustly upregulated the pRB (Ser780)-positive cells compared to negative-control siRNA-treated cells (Fig. 5e). These findings suggest that miR-425-5p may directly bind to the *PPP2CB* mRNA and suppress its expression, resulting in increased phosphorylation of RB and reduced cellular senescence. (Fig. 5f).

## Discussion

The results of the study clearly demonstrate that miR-425-5p mitigates cellular senescence following drug-induced senescence induction. Our findings raise the possibility that miR-425-5p may have therapeutic potential as a senomorphic agent for limiting the accumulation of therapy-induced senescent cells. Chemotherapy by drugs such as methotrexate or Doxorubicin induced senescence is increasingly recognized as a contributor to accelerated aging, tissue dysfunction, and chronic inflammation in cancer survivors through the persistent production of SASP factors^20^. Recent studies have demonstrated that modulation of specific miRNAs can mitigate these detrimental effects. For example, restoration of miR-199a-3p reduced doxorubicin-induced cardiac senescence and SASP production, while miR-7 suppressed therapy-induced senescence and restored gemcitabine sensitivity in pancreatic cancer cells^66,67^. Similar to these observations, miR-425-5p overexpression attenuated senescence, reduced SASP factor expression, and decreased DNA damage signaling following MTX treatment. These findings suggest that miR-425-5p may not only limit the long-term adverse consequences of therapy induced senescence but could also help prevent the establishment of persistent senescent cell populations that contribute to chronic inflammation and treatment resistance. Although future studies in primary cells and *in vivo* models will be required, our results identify miR-425-5p as a potential miRNA-based serotherapeutic candidate for reducing the burden of therapy-induced senescence.

In addition to attenuating therapy-induced senescence, miR-425-5p delayed the onset of replicative senescence in WI-38 fibroblasts, suggesting that its effects are not limited to acute stress responses. Replicative senescence is considered a physiologically relevant model of cellular aging driven by cumulative replication-associated damage. The delayed accumulation of senescent cells and enhanced proliferative capacity in miR-425-5p-overexpressing cells, supports a role for miR-425-5p in maintaining cellular homeostasis during cellular aging. Importantly, the ability of miR-425-5p to modulate both therapy-induced and replicative senescence suggests that it may regulate core senescence pathways that are shared across distinct senescence-inducing stimuli.

Interestingly, our data suggest that the anti-senescent effects of miR-425-5p may be dose dependent. While moderate overexpression of miR-425-5p attenuated senescence-associated phenotypes, higher levels failed to provide additional benefit and instead were associated with increased apoptosis. Such biphasic responses have been reported for other miRNAs and likely reflect the complex and highly interconnected nature of miRNA-mediated gene regulation. A notable example is miR-96, which exhibits opposing effects on autophagy depending on its expression level. Under hypoxic conditions, moderate increases in miR-96 promote autophagy through repression of *MTOR*; however, excessive miR-96 expression inhibits autophagy by targeting *ATG7*^68^. Similarly, studies of *let-7* family members and the miR-17-92 cluster have demonstrated nonlinear, dose-dependent target selection, whereby distinct targets are preferentially regulated at different miRNA concentrations, resulting in differential effects on cellular functions including viability^69^. These observations support the concept that miRNAs do not function as simple on/off regulators but rather as quantitative modulators of complex regulatory networks. Given that cellular senescence is governed by interconnected pathways controlling cell cycle arrest, DNA damage responses, apoptosis, autophagy, and inflammatory signaling, it is plausible that moderate miR-425-5p expression selectively represses senescence-promoting targets such as *PPP2CB* and components of the TGF-β pathway, whereas excessive expression may engage additional targets involved in cell survival and stress adaptation. Future studies defining the complete dose-response relationship and target repertoire of miR-425-5p will be important for understanding its role in senescence regulation and for determining the optimal therapeutic window for miR-425-based interventions.

## Acknowledgements

This work was supported by grants to FS from the NIH (5 R01AG082093, 5-R35 CA232105).

## Author Contributions

LM: Conceptualization, Investigation, Methodology, Project administration, Supervision, Writing-original draft. FS: Conceptualization, Funding acquisition, Project administration, Resources, Supervision, Writing-original draft. SH: Investigation, Methodology. JDL: Data analysis.

## Data Availability statement

The data that support the findings of this study are will be available upon request.

## Experimental Procedures

### Cell Culture

HEK293T (CRL-3216™), A549 (CCL-185^™)^, and WI-38 (CCL-75™), cells were purchased from the American Type Culture Collections (ATCC) and were cultured as per their recommendations. HEK293T cells were cultured in 1X Dulbecco’s Modified Eagle Medium (DMEM) (Gibco) supplemented with 10% Fetal Bovine Serum (FBS) and 1% penicillin-streptomycin (PS) (Gibco). A549 cells were cultured in 1X Roswell Park Memorial Institute (RPMI) 1640 Media (Gibco) supplemented with 10% FBS and 1% PS. WI-38 cells are cells isolated from the lung tissue of a 3-month-old, female, embryo and were cultured using ATCC-formulated Eagle’s Minimum Essential Medium supplemented with 10% FBS and 1% PS. All cell lines were cultured at 37°C and 5% CO2.

### miRNA mimic, inhibitor, and siRNA Transfections

A549 cells (2 × 10⁵ cells/well) were seeded in 6-well plates 24 h prior to transfection. miR-425-5p mimic (Assay ID: MC11575; Catalog #4464066), miR-425-5p inhibitor (Assay ID: MH11575; Catalog #4464084), and PPP2CB-targeting siRNA (Assay ID: s10960; Catalog #4390824) were purchased from Thermo Fisher Scientific.

Unless otherwise indicated, cells were transfected with 10 pmol of miR-425-5p mimic or the corresponding negative control mimic using Lipofectamine™ 3000 (Thermo Fisher Scientific) according to the manufacturer’s forward-transfection protocol. For miR-425-5p inhibition experiments, A549 cells were transfected with 30 pmol of miR-425-5p inhibitor or the corresponding negative control inhibitor using Lipofectamine™ RNAiMAX (Thermo Fisher Scientific). For PPP2CB knockdown experiments, cells were transfected with 30 pmol of validated PPP2CB siRNA or negative control siRNA using Lipofectamine™ RNAiMAX (Thermo Fisher Scientific).

### Lentiviral overexpression of miR-425-5p

A lentiviral vector expressing miR-425-5p under the control of the CMV promoter (Catalog #MH40628) and the corresponding negative control vector were purchased from ABM Good. Lentiviral particles were generated by co-transfecting the miRNA expression vectors with the packaging plasmid psPAX2 and the envelope plasmid pMD2.G (VSV-G) into HEK293T cells. Viral supernatants were collected and used for transduction experiments.

WI-38 cells at population doubling level (PDL) 35 were transduced with lentiviral particles for 48 hours, followed by puromycin selection to establish stable miR-425-5p-overexpressing and control cell populations. These cells were subsequently used for senescence and molecular analyses.

### SA-β -galactosidase assay

Cellular senescence was assessed using the Senescence β-Galactosidase Staining Kit (Cell Signaling Technology, #9860) according to the manufacturer’s instructions. Briefly, cells were seeded in 6-well plates and cultured under the indicated experimental conditions. Following treatment, cells were washed once with phosphate-buffered saline (PBS) and fixed for 10–15 min at room temperature using the fixative solution provided in the kit. After fixation, cells were washed twice with PBS and incubated with freshly prepared β-galactosidase staining solution (PH=6) at 37°C in a dry incubator without CO₂ overnight.

Senescent cells were identified by the development of a blue precipitate and imaged using a bright-field microscope. For quantification, at least five randomly selected fields per well were captured, and the percentage of SA-β-gal-positive cells was calculated by dividing the number of blue-stained cells by the total number of cells. A minimum of 100 cells per condition were analyzed. Experiments were performed in at least three independent biological replicates.

### Proliferation assays

Cells were plated in triplicates at 10K cells/well in a 96-well plate, including media only control wells. For proliferations, cells were plated after 3 days after washing the methotrexate drug. CyQuant MTT assay was performed the next day after plating as per manufacturers protocol. Briefly, 10μl of 12mM MTT was added to the cells after changing the media with 100μl of fresh media. Cells were incubated at 37°C in the dark for 3 hours after which 75 ul media was removed and 50 ul of DMSO was added. Plates were incubated at 37°C with optical shaking for 10 mins and then absorbance was measured at 560 nm using GloMax Explorer (Promega). Samples were normalized to cells treated with Neg-control and no drug treatment.

### Apoptosis Assay

Apoptosis was assessed using the RealTime-Glo™ Annexin V Apoptosis Assay (Promega, Cat. # JA1000) according to the manufacturer’s instructions. A549 cells were treated with increasing conc of miR-425-5p mimic for 36 hours. Cells were washed off the miRNA mimic and seeded in 96-well plates at 10,000 cells per well in 100 µL complete growth medium and allowed to adhere overnight. The following day, 100 µL of 2X Annexin V detection reagent was added directly to each well, resulting in a final 1X reagent concentration. Annexin V-dependent luminescence was measured after 30 min using a multimode plate reader.

### RNA isolation, cDNA preparation and qPCR

Total RNA was isolated from cells using Trizol and Pure Link RNA mini kit (Invitrogen Catalog # 12183018A) and concentration was measured using the NanoDrop Spectrophotometer (ND-1000 Spectrophotometer). miRNA expression levels were determined by quantitative RT-PCR (qRTPCR) using TaqMan miRNA Assays (Applied Biosystems) Assays specific to mature miRNAs were performed as per the manufacturer’s published protocols. miRNA expression levels were normalized to endogenous control U6. The expression of miR-425-5p targets, *CDKN1A*, and cytokines mRNA was assayed using SYBR Green I according to manufacturer protocols (Roche).

The expression levels were normalized to mRNA levels of the GAPDH. qPCR primers were designed to span at least one exon-exon boundary.

### RNAseq and data analysis

At the end of the incubation period, cells were harvested, and total RNA was isolated as described above. RNA quality and integrity were assessed using the Agilent 2100 Bioanalyzer and RNA 6000 Nano Kit (Agilent Technologies, Santa Clara, CA, USA). cDNA library preparation and RNA sequencing were performed by the Harvard Biopolymer Core Facility. Libraries were sequenced on an Illumina NextSeq 500 platform using 75-bp paired-end sequencing. Raw fastq reads were aligned with salmon quantification to the ensemble v104 reference transcriptome. Gene counts were imported into DESeq2, and differential expression analysis was performed using donor as a covariate. Raw RNA-seq datasets are available upon reasonable request.

### Protein Isolation and Immunoblot

Whole-cell lysates were prepared in radioimmuno-precipitation assay (RIPA) buffer (Invitrogen) supplemented with 1X protease inhibitor cocktail, and the DNA was sheared by passing through a 27.5-gauge needle. Lysates were centrifuged for 30 min at 14,000 rpm at 4 °C to clear cell debris and then total protein concentrations were determined using Pierce BCA Protein Assay Kit (Thermo Scientific). Equal mass of each sample was denatured in 4X Laemmli sample buffer with β-mercaptoethanol by boiling at 95°C for 10 minutes. Samples were loaded onto a NuPAGE 4-12% Bis-Tris gel and run in MOPS buffer. Gels were transferred onto nitrocellulose membrane via wet overnight transfer at 30V, blocked with blocking buffer (5% wt/vol milk or BSA in 1X TBST) for 1 h at room temperature (RT). Membranes were incubated with primary antibodies with gentle agitation overnight at 4 °C and then incubated with horseradish peroxidase (HRP)-linked secondary antibodies for 1h at RT. Membranes were imaged using SuperSignal West Femto substrate (Thermo Fisher Scientific) on Amersham Imager 600. Relative protein levels were quantified using the Fiji software compared to β-actin. All antibodies used are listed here:

| <u>Antibody</u> | <u>Source</u> | <u>Catalog #</u> | <u>Dilution used</u> |
| --- | --- | --- | --- |
| γ-H2AX | Abcam | ab81299 | 1:1000 in 5% BSA |
| H2AX | CST | 2595S | 1:1000 in 5% milk |
| ATM (Y170) | Abcam | Ab32420 | 1:1000 in 5% milk |
| pATM (s1981) | Abcam | Ab81292 | 1:1000 in 5% BSA |
| RB | CST | 9309S | 1:1000 in 5% milk |
| pRB (s780) | CST | 9307T | 1:1000 in 5% BSA |
| P53 | CST | 9282S | 1:1000 in 5% BSA |
| Cyclin D1(SP4) | Abcam | Ab16663 | 1:1000 in 5% milk |
| $\beta$ -actin | SCBT | Sc47778 | 1:1000 in 5% milk |

## Supplementary information

### Supplementary Figure Legends

**Figure S1:**
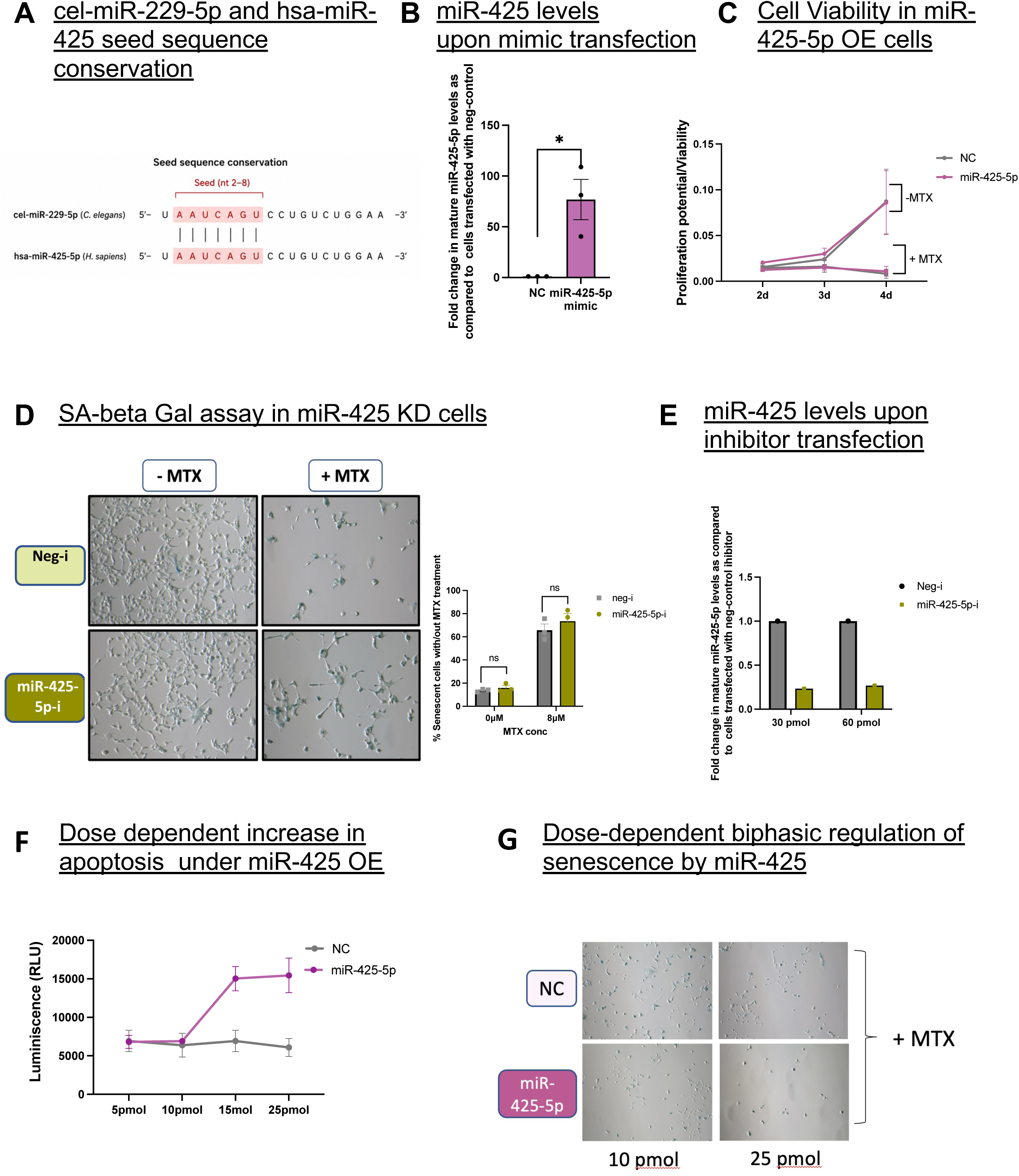
A) Multiple sequence alignment indicates 100% seed sequence conservation between cel-miR-229-5p and hsa-miR-425-5p. B) The graph represents relative expression of mature miR-425-5p in A549 cells treated with 10pmol of miR-425-5p mimic as compared to cells treated with 10pmol of Negative control mimic (NC). The cells were collected at the same time point as the other senescence related assays were performed. * indicates p<0.05, as. determined by a student’s t-test. C) Quantification of cell proliferation in negative control and miR-425-5p-overexpressing A549 cells after MTX or no drug treatment, as measured by the CyQUANT MTT assay after 48 (2d), 72 (3d) and 96 (4d) hours post MTX recovery. miR-425-5p overexpression slightly increased proliferation under untreated conditions compared with negative control cells. D) Representative images and quantification of the percentage of senescent cells following a 72-hour recovery period after MTX treatment or basal conditions in A549 cells transfected with either a negative control inhibitor or miR-425-5p inhibitor. Quantification demonstrate a marked increase in the percentage of senescent (blue) cells following miR-425-5p inhibition, indicating that loss of miR-425-5p exacerbates MTX-induced cellular senescence. E) Graph showing the relative expression of mature miR-425-5p in A549 cells transfected with 30 pmol or 60 pmol of miR-425-5p inhibitor compared with cells transfected with the corresponding amounts of negative control inhibitor. F) Quantification of apoptosis in negative control and miR-425-5p-overexpressing A549 cells under untreated and senescence-inducing conditions, as measured by the luminescence generated in the Annexin V apoptosis assay. G) Representative images of senescent A549 cells transfected with increasing concentrations of miR-425-5p mimic prior to senescence induction, indicating no further reduction in senescence at higher miR-425-5p mimic concentrations.

**Figure S2:**
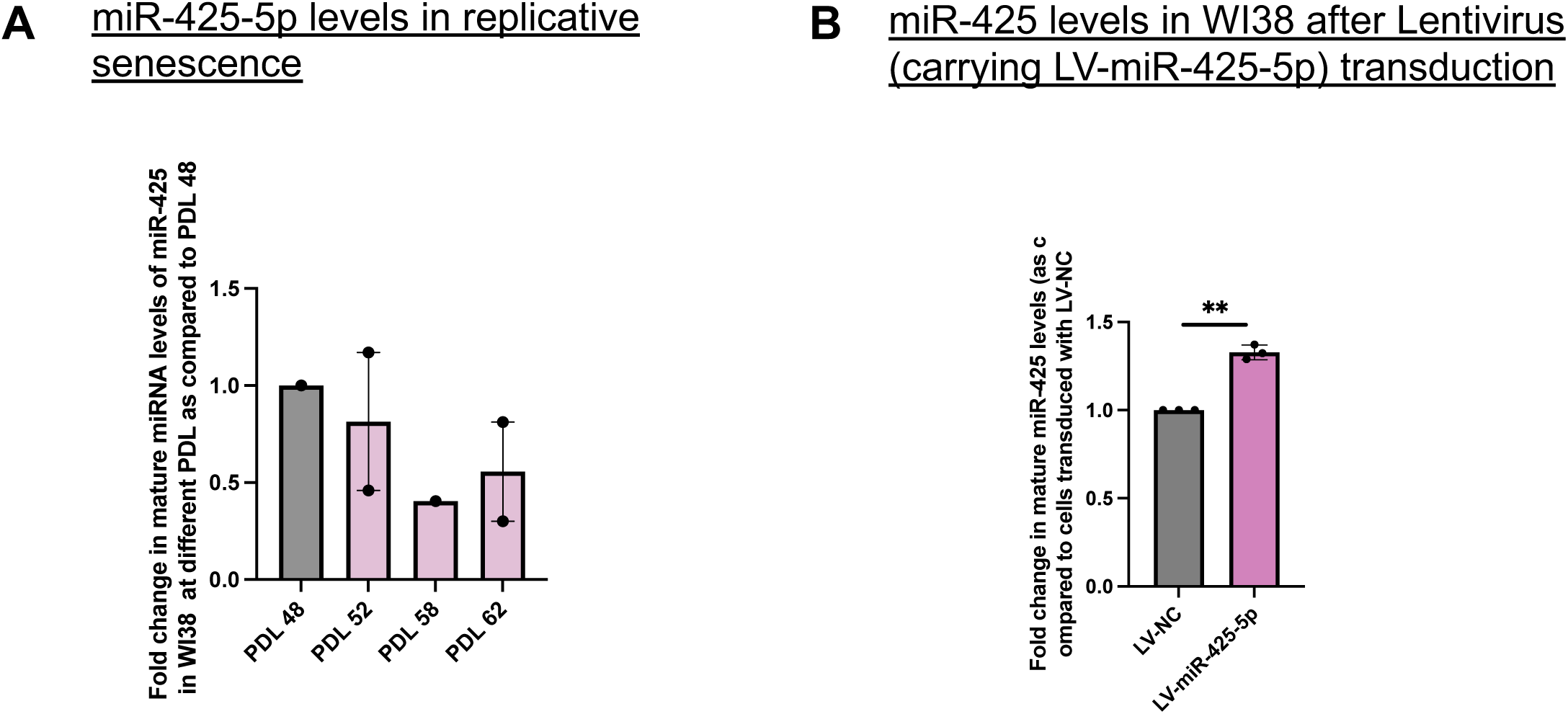
A) Graph showing the relative expression of mature miR-425-5p at the indicated population doubling levels (PDLs), normalized to PDL 44. B) Graph showing the relative expression of mature miR-425-5p in WI-38 cells transduced with a miR-425-5p-overexpressing lentiviral vector compared with cells transduced with a negative control lentiviral vector. Error bars represent SEM from three independent biological replicates. * indicates p < 0.05, as determined by Student’s t-test.

**Figure S3:**
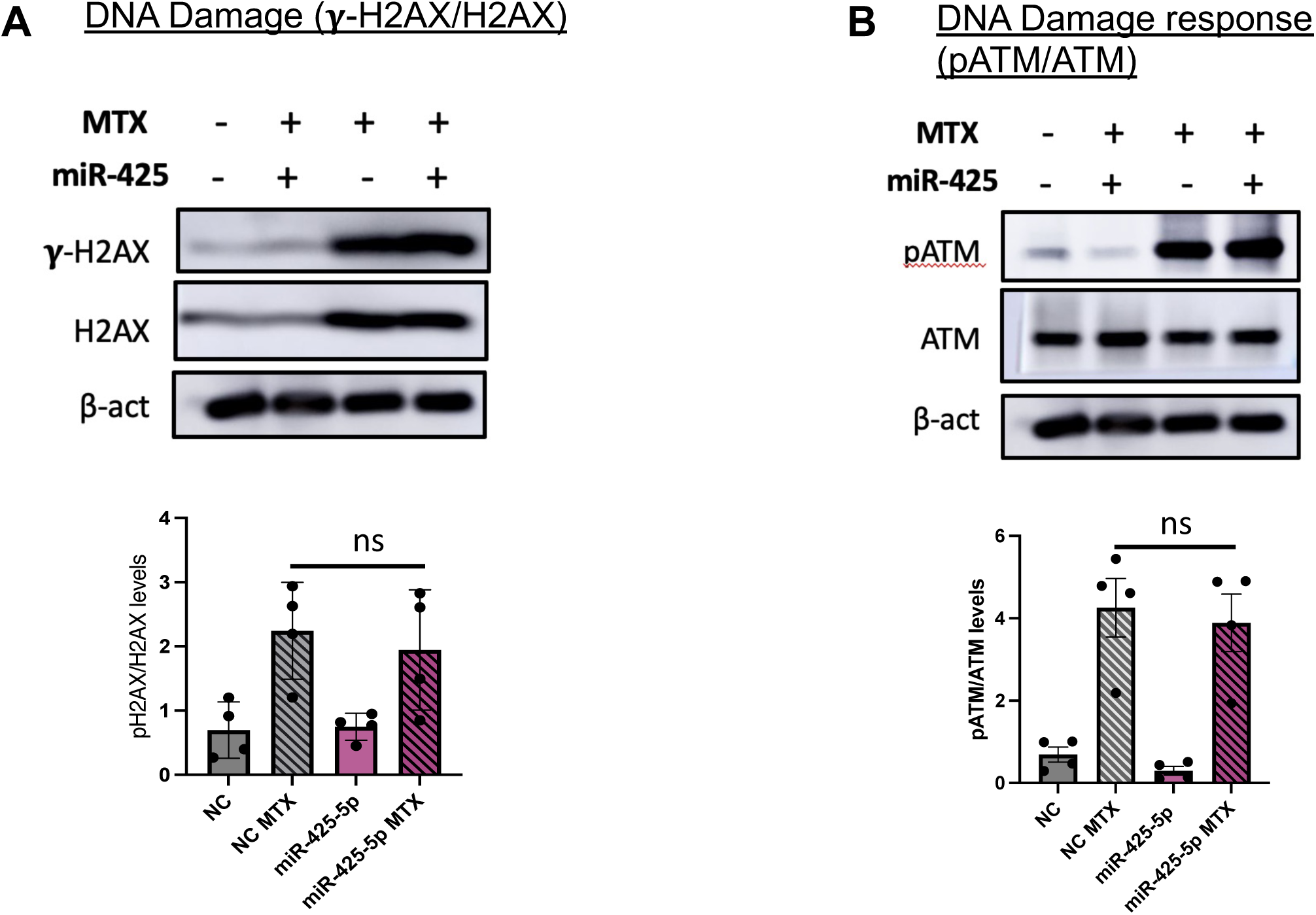
A) Representative western blots and corresponding quantification of γ-H2AX (S139), total H2AX, and β-actin levels in negative control and miR-425-5p-overexpressing A549 cells under untreated and MTX-induced senescence conditions. The graph on the right shows quantification of γ-H2AX levels normalized to total H2AX levels, with total H2AX levels normalized to the loading control β-actin. Data represent three independent biological replicates. *indicates p<0.05, as. determined by a student’s t-test. (B) Representative western blots and corresponding quantification of pATM (S1981), total ATM, and β-actin levels in negative control and miR-425-5p-overexpressing A549 cells under untreated and MTX-induced senescence conditions. The graph on the right shows quantification of pATM levels normalized to total ATM levels, with total ATM levels normalized to the loading control β-actin. Data represent three independent biological replicates. *indicates p<0.05, as. determined by a student’s t-test.

**Figure S4:**
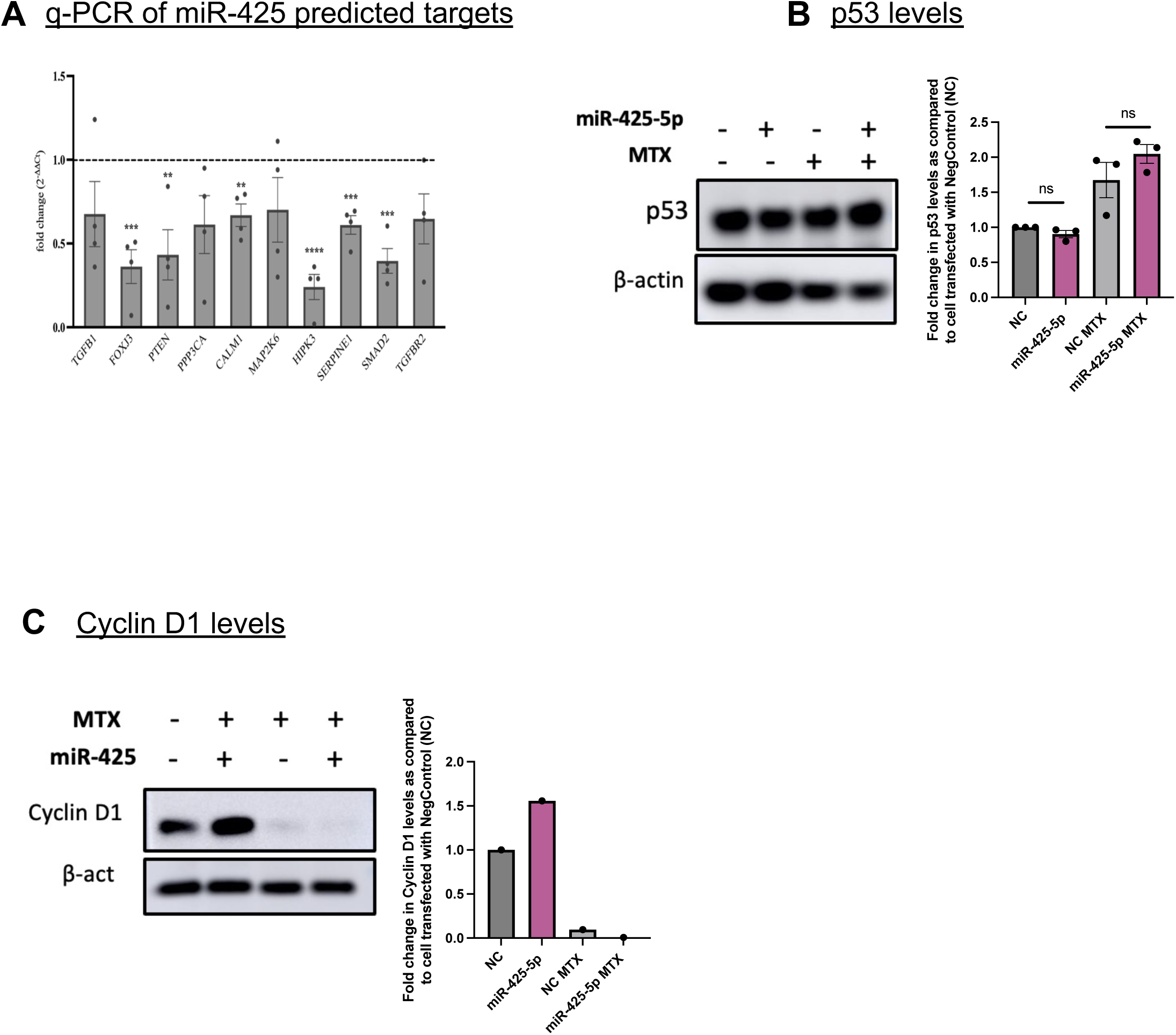
A) Graph represents fold changes in mRNA levels of miR-425-5p predicted targets in neg-control A549 cells treated with MTX, miR-425-5p OE cells, untreated and treated with MTX, as compared untreated negative control cells (NC). Error bars show SEM, *indicates p<0.05, as. determined by a student’s t-test. B) Representative western blots and corresponding quantification of p53 and β-actin levels in negative control and miR-425-5p-overexpressing A549 cells under untreated and MTX-induced senescence conditions. The graph on the right shows quantification of p53 levels normalized to the loading control β-actin. Data represent three independent biological replicates. *indicates p<0.05, as. determined by a student’s t-test. (C) Representative western blots and corresponding quantification of Cyclin D1 and β-actin levels in negative control and miR-425-5p-overexpressing A549 cells under untreated and MTX-induced senescence conditions. The graph on the right shows quantification of Cyclin D1 levels normalized to the loading control β-actin.

**Figure S5:**
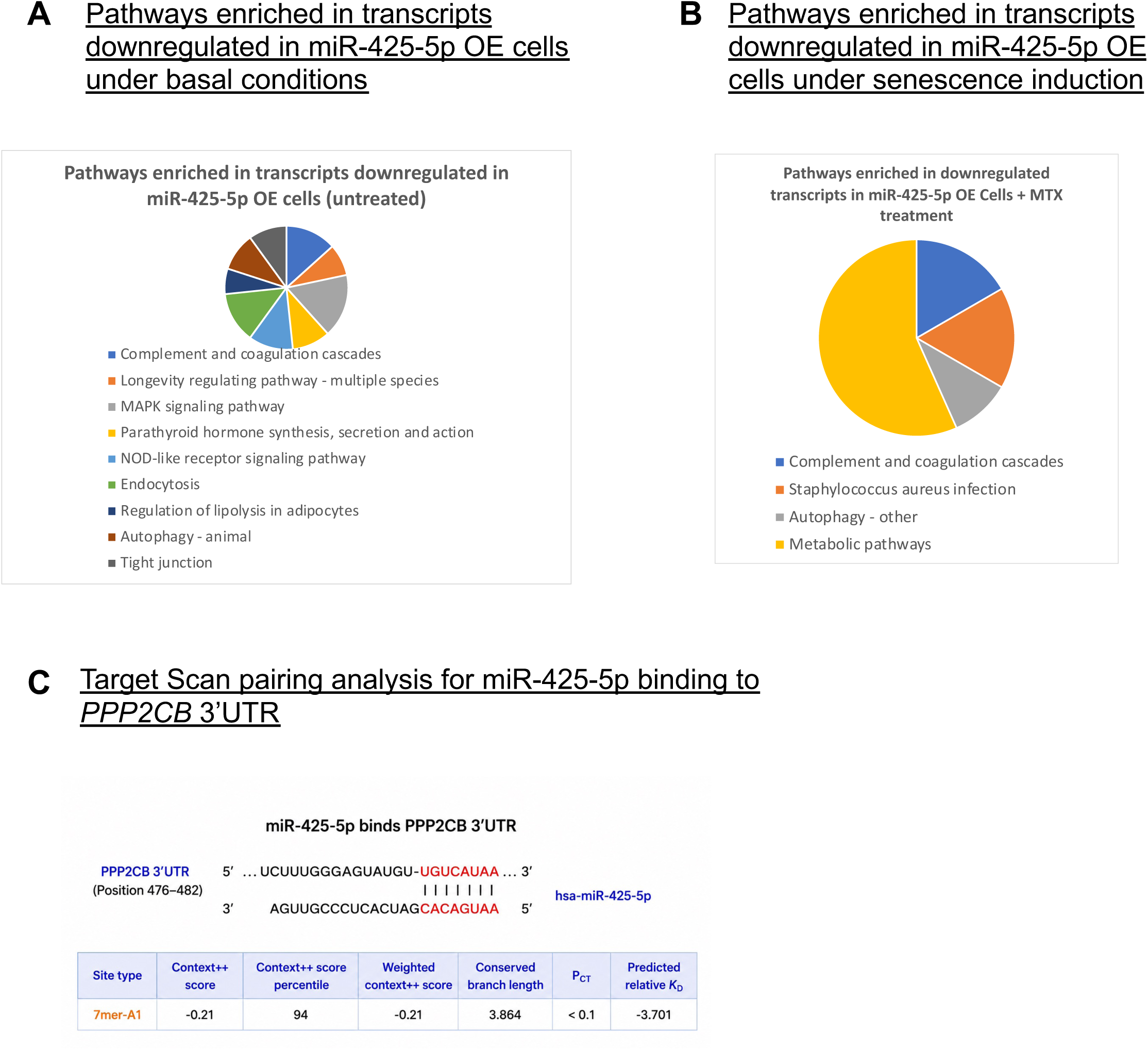
A) GO and KEGG pathway enrichment analysis of transcripts downregulated in miR-425-5p-overexpressing A549 cells compared with negative control cells under basal conditions. Enriched pathways (FDR < 0.1) were associated with inflammatory signaling, longevity regulation, MAPK signaling, NOD-like receptor signaling, autophagy, endocytosis, lipid metabolism, and cell junction organization. B) GO and KEGG pathway enrichment analysis of transcripts downregulated in miR-425-5p-overexpressing A549 cells compared with negative control cells following MTX-induced senescenc0e. Enriched pathways (FDR < 0.1) were associated with complement and coagulation cascades, host defense and inflammatory responses, autophagy, and metabolic regulation. C) TargetScan analysis showing a conserved miR-425-5p binding site within the PPP2CB 3′ untranslated region (3′UTR). The predicted interaction corresponds to a 7mer-A1 seed match with a Context++ score of −0.21 (94th percentile) and a predicted relative K_D of −3.701.

